# Aerobic bacteria as *Escherichia coli* can survive in ESwab^TM^ medium after a 3 month-freezing at −80°C but not after multiple thawing

**DOI:** 10.1101/537647

**Authors:** Rindala Saliba, Jean Ralph Zahar, Etienne Carbonnelle, Mathilde Lescat

## Abstract

Pretesting of procedures for specimen conservation should be part of preliminary studies for trials especially when quantification of bacteria must be performed in a second time. Quantitative epidemiological studies of multidrug resistant organisms sampled from rectal swabs could be then particularly favored. The aim of this preliminary study, was to evaluate the performance of the flocked swab with liquefied Amies transport medium ESwab™, for the survival evaluation of aerobic bacteria, from rectal swab sampling according to the number of freezing at −80°C and thawing cycles and the time of freezing. We first observed that quantification was not reliable after F/T cycles whereas a unique freezing could be performed especially when studying *E. coli* isolates. The second experiment allowed us to observe that this stability could be obtained until 3 month-freezing. Our study represents a preliminary study, confirming the utility of ESwab™ in microbiological diagnostics and research studies, not only for molecular bacterial tests, but also, for the maintenance of bacterial viability in clinical specimens.

## Introduction

Understand the bases of emergence of multidrug resistant organisms (MDRO) has become an important and real challenge those last decades. *Enterobacteriaceae* are the most frequent MDRO spreading worlwide and are usually carried by patients in microbiota before being involved in pathologies (and isolated in clinical samples). Among this group, *Escherichia coli* can be considered as one of the most important because it is the most performant human gut commensal and become, depending on factors related to the host or the strain, a formidable pathogen (1). Besides, nowadays extended βlactamase (ESBL) producing isolates belong mainly to this species in both hospitals and community and constitute a major public health concern worldwide (2). Moreover, a high relative abundance of ESBL *E. coli* has been associated with longer fecal carriage time and a higher risk of infection (3). Many *in vivo* and *in vitro* studies try to identify the basis of commensalism which is natural the behavior of *E. coli* isolates whether it is resistant or not (4). The other MDRO belong mainly to the ESKAPE group (*Enterococcus faecium, Staphylococcus aureus, Klebsiella pneumoniae, Acinetobacter baumannii, Pseudomonas aeruginosa*, and *Enterobacter* species) (5). Likewise, those species are also opportunistic pathogens and quantitative studies of the prevalence, bacterial load and diversity of these species as well as *E. coli* in their natural environment, which is the digestive gut, are essential (3, 6, 7).

Several steps in the analyses of bacteria from digestive gut are crucials and are directly related to a good diagnostic performance in the Clinical Microbiology laboratory. For example, in molecular methods, it is well known that DNA extraction or library preparation are key steps that could influence the quality of results. Besides, and whatever the method used (molecular or culturomics), it is essential that the sampling, transport and storage procedures do not alter the microbial composition (8). The best results are obtained when viability of micro-organisms is maintained, as well as, the relative proportions of all micro-organisms present in the clinical specimens and when liquid swab transport are used (9, 10). However, the studies evaluating the stability of the bacteria in those transport devices do not exceed days. The transport device, ESwab™ is the combination of a nylon-flocked swab with a transport maintenance medium, a modified liquid Amie’s medium. ESwab™ is a swab systeme presenting advantages compared to other, including enhanced uptake and release of bacteria due the flocked nylon, the bacterial survival in the medium and an easy to use system. Swab specimens for bacterial investigations collected using ESwab™, that can’t immediately delivered to the laboratory, should be refrigerated at 4-8°C or stored at room temperature (20-25°C) and processed within 48 hours maximum of collection. According to the manufacturer, ESwab™ could be freezable at −20°C; however, after thawing, only molecular bacterial tests should be performed (13). Nowadays, numerous labs use ESwab™ system to sample the patients and evaluating this device in quantitative studies could be useful for hospitals in which patients are sample with it.

So far quantitative studies of *E. coli* in the feces have been performed using fresh feces or rectal swabs rapidly seeded or placed in Brain Heart infusion (BHI) with glycerol and placed at −80°C (3, 6, 7). In prospective studies, these procedures are time consuming and can not be applied in numerous labs. Quantitative studies with other bacteria are very rare, based on total microbiota analyses or absent and these studies are needed (11, 12). To have the opportunity to freeze directly available devices could be useful in many labs and increase the feasibility of these experiments.

Pretesting of procedures for specimen collection, transport and conservation should be part of preliminary studies for trials. The aim of this preliminary study, was (i) to evaluate the survival of aerobic bacteria from rectal swab placed in the transport medium ESwab™ according to the number of freezing at −80°C and thawing cycles (F/T cycles) and (ii) the survival of *E. coli* isolates from rectal swab placed in the transport medium ESwab™ during a long period (3 months).

## Materials and Methods

The survival of aerobic bacteria in the ESwab™ (Copen Diagnostics, Italy), devices was investigated by quantification of aerobic bacteriain two experimentations. Both experiments were realized using samples of clinical rectal ESwab™, chosen randomly, in the laboratory of Microbiology of Jean Verdier Hospital, Bondy, France. In the first experimentation, 9 samples were treated during a period of 3 weeks, by two methods A and B of F/T cycles for comparison after an initial quantification of each aerobic bacterium.

In the second experimentation 4 samples were treated during 3 months by the method B for comparison after an initial quantification of *E. coli* isolates.

In the method A, an aliquot of 400 µL of the initial medium suspension of the ESwab™ sample was stored at −80°C and defreezed and refreeze (F/T cycles) at each step (1 week, 2 weeks and 3 weeks) In the method B, three aliquots of 100 µL each were frozen at −80°C and used for quantification at each step (1 week/month, 2 weeks/months and 3 weeks/months, according to the experimentation). Each aerobic bacterium quantification was performed using 100 µL of the medium suspension contained in the ESwab™. The suspension was first serially 10-fold-diluted. Then, a 100-µL sample of each dilution was inoculated on an UriSelect4 plate (Bio-Rad, Marnes, la coquette, France) and spread over the entire surface using a sterile L-spatula. Plates were incubated for 18-24h at 37°C. The quantification of each aerobic bacterium was obtained by performing each aerobic bacteria colony count on three dilution plates. The relative quantification for each point was obtained by dividing the result of each aerobic bacterium quantification at each time (1 week/month, 2 weeks/months and 3 weeks/months according to the experimentation) by the initial result of each aerobic bacterium quantification at time 0 (TO).

For all experiments a first study of one type of colony of bacteria was realized at TO allowing the identification at the species level using the Microflex bench-top Matrix Assisted Laser Desorption Ionisation-Time of Flight (MALDI-TOF) mass spectrometer (Brücker, Champs-sur-Marne, France). Then, we decided to interpret the identification of all other bacteria with the same size, colour and aspect of the colonies after a first identification by MALDI-TOF.

All statistics were computed performed using R software (R Development Core Team, 2009, Vienna Austria) and statistical significance was determined at a p-value of less than 0.05.

## Results

In the first experiment, the survival of each aerobic bacterium in the ESwab™ devices was evaluated, by calculating a relative quantification ratio, based on each aerobic bacterium colony count, then, by determining the relative ratio using the two methods A and B. Among the 9 samples, 6 were positive for *E. coli* detection, two for *K. pneumonia* and 8 for *Enterococcus species* at TO. When detected all aerobic bacterium quantification were comprised between 2.0 10^4^ and 1.5 10^7^ colony forming unit (CFU)/mL of ESwab™ liquid (Table1). When we observed aerobic bacterium relative quantification across the time, variations were observed between the two methods and the different bacteria. The relative quantification of both *Enterobacteriaceae* across the time was similar with a drastic decreased quantification (no significant difference using t-test) when samples were subjected to F/T cycles (method A). On the contrary, in the method B, quantifications remained stable for both *Enterobacteriaceae* with a significant higher resistance of *E. coli* without inactivation (ratio= 1 all along the experiment) whereas *K. pneumonia* quantifications were recovered at 60% (p=5.7 10^-5^). *Enterococcus species* remained stable at a relative quantification at 80% when aliquoted at the beginning and slightly decreased at 40% when freezed and defreezed. When we compared *E. coli* and *Enterococcus species* inactivations, we observed that *E. coli* were more resistant to long freezing states without thawing (method B), whereas *Enterococcus species* were more resistant to F/T cyles (method A) (p=0.02 and 0.04, respectively).

In the second experiment, we performed the freezing during three months of 4 other Eswab™ sample aliquots containing exclusively *E. coli* isolates to determine specifically for this species during a longer period if the stability could be observed. Indeed, we observed a complete stability of the quantification of *E. coli* isolates at a mean value of 2 10^6^ CFU/mL during the three months.

## Discussion and conclusion

In this study, we evaluated the survival of aerobic bacteria from rectal ESwab™ according F/T cycles and then the survival of *E. coli* isolates from rectal swab placed in the transport medium Eswab™ during a long period (3 months).

Our first results indicated that all aerobic bacterium initial quantification were comprised between 2.0 10^4^ and 1.5 10^7^ colony forming unit (CFU)/mL of Eswab™ liquid (Table1) which is decreased compared to 16S quantification analyses of aerobic bacteria comprised in the total microbiota obtained when stools are collected instead of rectal samples (14). This could be explained by the fact that this quantification was performed after a per mL preliminary dilution of the feces in the Eswab™ liquid. In fact these quantifications are consistant with *E. coli* quantification, (from the most frequent aerobe isolated from human feces) which is about 10^8^ colony forming unit (CFU) per gram of feces (14). It was not possible to weight the feces retrieved on samples due to the fact that analyses were performed from patient samples. And this should be taken into account in quantification analyses.

**Table 1:** Initial quantification of aerobic bacteria in the 9 ESwab™ samples. The numbers correspond to the numbers of colony forming unit per mL of ESwab™ liquid.

|  | *E. coli* | *K. pneumoniae* | *Enterococcus species* |
| --- | --- | --- | --- |
| **Sample 1** | 1.2 10<sup>5</sup> | ND | 1.1 10<sup>5</sup> |
| **Sample 2** | 5.5 10<sup>6</sup> | ND | 2.5 10<sup>5</sup> |
| **Sample 3** | 8.5 10<sup>6</sup> | ND | 1.2 10<sup>5</sup> |
| **Sample 4** | 2.0 10<sup>5</sup> | ND | 1.5 10<sup>7</sup> |
| **Sample 5** | ND | ND | 7.0 10<sup>6</sup> |
| **Sample 6** | ND | 7.5 10<sup>6</sup> | 2.0 10<sup>4</sup> |
| **Sample 7** | 2.0 10<sup>5</sup> | ND | 2.5 10<sup>6</sup> |
| **Sample 8** | ND | 3.0 10<sup>5</sup> | 6.0 10<sup>6</sup> |
| **Sample 9** | 8.2 10<sup>5</sup> | ND | ND |

**Figure 1:**
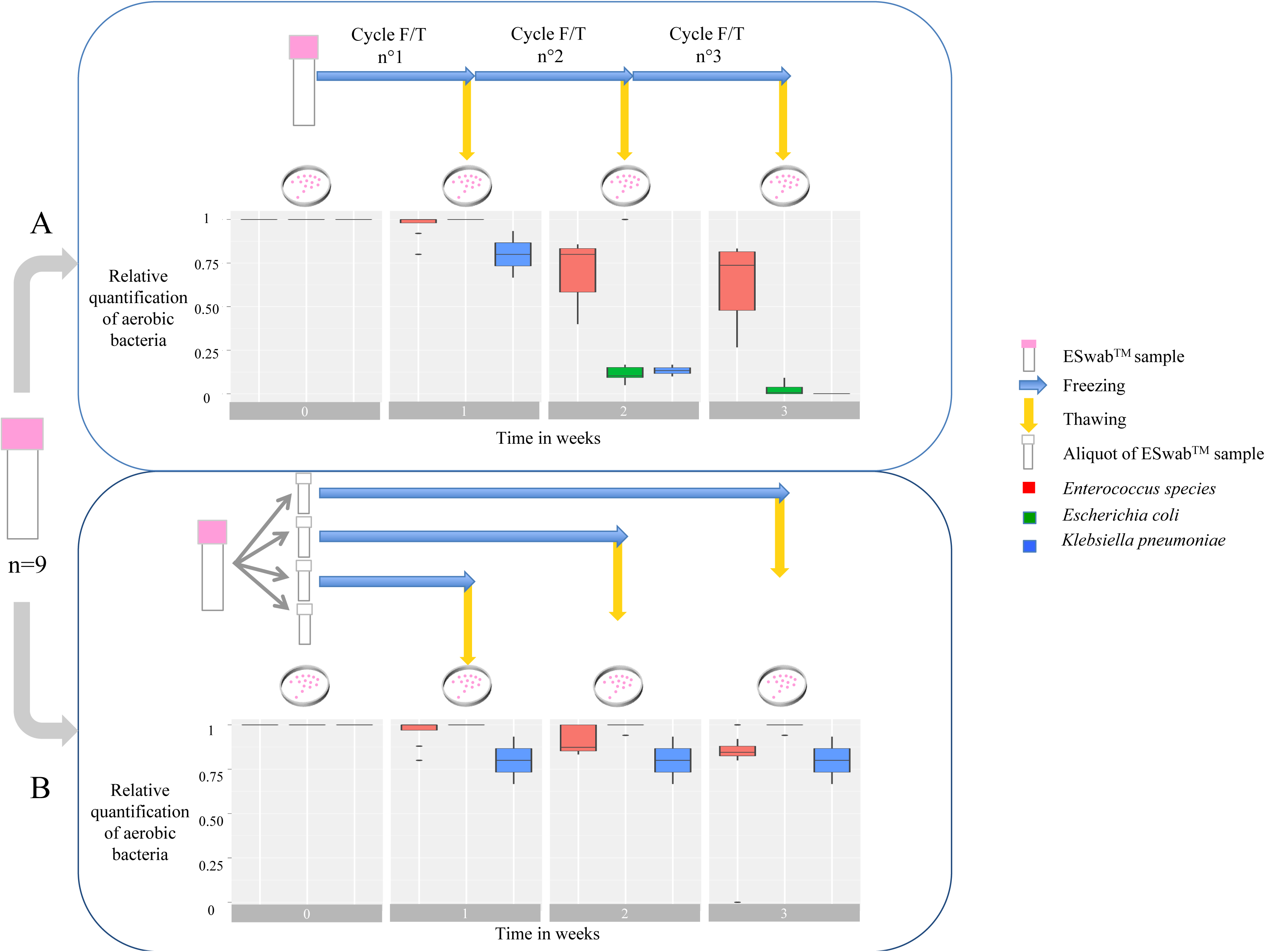
Boxplots of the relative quantification of aerobic bacteria in 9 ESwab™ samples during the first evaluation of 3 weeks, stored in 2 different methods A and B. The relative quantification was calculated, by dividing each aerobic bacterium quantification by the initial aerobic bacterium quantification (at time 0). The symbols are indicated. In the boxplot figure, when a line replace a boxplot means that there is no variations in the results of ratio of the isolate from the species.

Our findings suggest that the Eswab™, is able to preserve the initial quantity of *E. coli* when aliquoted and frozen at −80°C during minimum 3 months but F/T cycles altered significantly the initial amount of isolates. *Enterococcus species* isolates are retrieved after cycles of F/T whatever the number of cycles (maximum 3) but the stability is increased when aliquots are performed at the beginning (from 40% to 80%). We were not able to explain such discrepancies in bacteria families and only found the study of Gao *et al* who compared resistance of isolates of *E. coli* and *E. faecalis* freezed in sterile water in different temperatures. They also observed increased inactivation of all the isolates proportional to the number of F/T cycles with higher resistance of *E. faecalis*. They did not observe higher resistance of *E. coli*, but the freezing duration is not indicated in the material and method section and they used water as liquid of conservation (15).

The best way to obtain quality results for both molecular and culturomic methods is to use fresh feces from individuals as samples. This procedure can be easily performed when subject of a study are healthy individuals or laboratory animals. However, the collection of feces when subjects of studies are patients can be more difficult to manage. The use of rectal swabs followed by freezing seems the preferential procedure in the context of the difficulty of managing patient in the units and the samples in the lab. In fact, adequate strategies are required to limit bias due to shifts in microbial communities during sampling and storage. Rectal swabs are relatively simple samples to collect and are easily transported to the laboratory (16). They are routinely used in clinical settings to detect enteropathogens and multidrug resistant *Enterobacteriaceae* by culture analysi*s*. Knowing that some of them, like ESwab™, are also suitable for molecular analysis, they might be used to study the fecal microbiota composition. Moreover, studies have shown that rectal swabs are an acceptable and practical proxy alternative to stool, for the collection of fecal specimens and microbiota analysis (8, 16, 17). In some cases the determination of relative density of aerobic bacteria is of particular interest in some specific clinical questions whether the study is a one point or longitudinal follow up. For example, fecal density of ESBL *E. coli* has been observed as an important risk factor in ESBL infection (3). Methodology of transport and conservation of fecal samples is then crucial before analysis as well as culture conditions (mainly media).

Although fecal swabs are often used in the clinical setting, because of being the most user friendly method to obtain fecal samples, to our best knowledge, this is the first study that investigates the effect of F/T cyles and long freezing on the viability of aerobic bacteria *as E. col*i, in the ESwab™. Our study represents a preliminary study, confirming the utility of ESwab™ in microbiological diagnostics and research studies, not only for molecular bacterial tests, but also, for the maintenance of bacterial viability in clinical specimens allowing combined quantitative study of aerobic bacteria (*E. coli*) with total microbiota by molecular analyses. Optimal results are obtained when ESwab™ initial medium suspension is aliquoted and stored at −80°C. Then thawing of aliquots must be performed once for each. However due to the limited number of samples observed and especially for *K. pneumoniae*, our study should be completed, on a larger scale, by testing more samples.

